# Identification of robust cellular programs using reproducible LDA that impact sex-specific disease progression in different genotypes of a mouse model of AD

**DOI:** 10.1101/2024.02.26.582178

**Authors:** Narges Rezaie, Elisabeth Rebboah, Brian A. Williams, Heidi Yahan Liang, Fairlie Reese, Gabriela Balderrama-Gutierrez, Louise A. Dionne, Laura Reinholdt, Diane Trout, Barbara J. Wold, Ali Mortazavi

**Affiliations:** Department of Developmental and Cell Biology, University of California, Irvine, CA, USA; Center for Complex Biological Systems, University of California, Irvine, CA, USA; Division of Biology, California Institute of Technology, Pasadena, CA, USA; The Jackson Laboratory, Bar Harbor, ME, USA

## Abstract

The gene expression profiles of distinct cell types reflect complex genomic interactions among multiple simultaneous biological processes within each cell that can be altered by disease progression as well as genetic background. The identification of these active cellular programs is an open challenge in the analysis of single-cell RNA-seq data. Latent Dirichlet Allocation (LDA) is a generative method used to identify recurring patterns in counts data, commonly referred to as topics that can be used to interpret the state of each cell. However, LDA’s interpretability is hindered by several key factors including the hyperparameter selection of the number of topics as well as the variability in topic definitions due to random initialization. We developed Topyfic, a Reproducible LDA (rLDA) package, to accurately infer the identity and activity of cellular programs in single-cell data, providing insights into the relative contributions of each program in individual cells. We apply Topyfic to brain single-cell and single-nucleus datasets of two 5xFAD mouse models of Alzheimer’s disease crossed with C57BL6/J or CAST/EiJ mice to identify distinct cell types and states in different cell types such as microglia. We find that 8-month 5xFAD/Cast F1 males show higher level of microglial activation than matching 5xFAD/BL6 F1 males, whereas female mice show similar levels of microglial activation. We show that regulatory genes such as TFs, microRNA host genes, and chromatin regulatory genes alone capture cell types and cell states. Our study highlights how topic modeling with a limited vocabulary of regulatory genes can identify gene expression programs in singlecell data in order to quantify similar and divergent cell states in distinct genotypes.

## Introduction

The different cell types constructing a tissue work together to carry out the functions of that tissue in response to various developmental and environmental cues by activating specific cellular programs. Single-cell and single-nucleus RNA sequencing (RNA-seq) enable the identification of cell types, subtypes, and cell states through their single-cell transcriptomes, enhancing our understanding of cellular phenotype heterogeneity and composition within complex tissues such as the brain(1–3). A common approach for cell type annotation relies primarily on unsupervised clustering methods(4), which partition cells based on the similarity of their gene expression patterns. This is followed by manual cell type assignment for each cluster based on differentially expressed markers from literature. The overall accuracy of this approach depends on both the clustering accuracy(5) and the prior knowledge of marker gene expression levels(6). For example, marker genes could be expressed in more than one cell type, complicating the annotation process. More importantly, this cluster-based approach assumes that cells can only be part of a single cluster, thereby averaging the cell-to-cell variability within that cluster.

Here, we focus on a key challenge of inferring complex cellular states and identities that are encoded by patterns of gene expression. We assume that each cell or nucleus engages in a limited number of cellular programs, and its observed transcriptome is determined by the sum of these active programs. To represent this, we leverage grade of membership (GoM) models(7, 8), allowing each cell to have partial membership in multiple cellular programs. One such model is Latent Dirichlet Allocation (LDA)(9), a probabilistic algorithm capable of inferring recurring combinations referred to as topics. LDA starts by randomly assigning topics to each word in a document, which in this case are cells(8). Due to this random initialization, different topic assignments and, consequently, different topic representations for each document may arise in repeated runs. As a result, the topics discovered by LDA can vary across different runs of the algorithm. To address this issue of topic variability, one common approach is to use a fixed random seed before running LDA, ensuring consistent random initialization across different runs. However, there is no guarantee that this fixed seed will produce the best topics, as some randomly defined topics might be stuck in local optima that lack biological significance.

Alzheimer’s disease (AD) is a progressive neurodegenerative disease characterized by memory loss(10). Microglia are the resident macrophages of the brain that mediate brain homeostasis by regulating immune function and promoting neuronal homeostasis and neuroprotection. To main-tain homeostasis, microglia damage or kill neurons with abnormal profiles(11, 12). However, not all the microglia in the brain behave identically. Single-cell RNA-seq studies have identified new microglial subtypes with unique transcriptional and functional characteristics, termed “disease-associated microglia” (DAM) in animal models of AD(13). DAMs are characterized by the upregulation of genes associated with late-onset AD, such as apolipoprotein E (*Apoe*), *Itgax, Csf1r*, and *Tyrobp*, whereas *Tmem119, Cd33*, and *Maf* are downregulated. However, in cluster-level analyses, these cells are typically treated either as part of distinct clusters or positioned on a pseudo time continuum between homeostasis and activation.

Here, we develop Topyfic, a Python package that (a) runs LDA multiple times with different random seeds, (b) aggregates similar topics across runs to compute reproducible topics, and (c) filters out low-participation topics. Using this strategy, we find reproducible topics, while simultaneously filtering out noisy, irreproducible topics. We apply Topyfic to single-nucleus and single-cell datasets generated by the ENCODE and MODEL-AD consortiums from mice with and without the 5xFAD transgene in either a C57BL/6J (MODEL-AD) or a C57BL/6J x CAST/EiJ background (ENCODE) to identify topics that are (a) detected in both genotypes with and without the transgene, and (b) microglia-specific. We then train additional topics with a subset of regulatory genes such as transcription factors and show that these regulatory topics that we recover also capture cell activation using regulatory genes alone.

## Results

### Reproducible LDA topics using Topyfic

Topyfic estimates the most likely number of topics while also maximizing the number of meaningful topics. The core idea of Topyfic is that topics that are found repeatedly across multiple LDA runs are more reliable than topics found in any single run, which could be suboptimal as the result of poor random initialization (Fig. 1A). Starting from a cell-by-gene expression matrix in h5ad format, Topyfic first trains a Latent Dirichlet Allocation (LDA) model on the provided dataset using different random seeds (see methods for a detailed technical description of Topyfic parameter settings). Topyfic then uses Leiden clustering(14) in gene-weight space to combine similar topics across individual runs to construct a consensus set of topics called the TopModel. Finally, Topyfic calculates cell-topic participation based on the TopModel and filters out any topic with user-defined low participation. Topyfic also includes several helper functions to analyze and visualize topics and topic participation in the training and testing datasets. Determining the number of topics is a challenging step for the application of LDA. We use two diagnostic metrics and visualizations to help guide this decision. First, we train our TopModels using a different starting number of topics, followed by pruning low-participation topics. In general, applying our consensus models to repeated runs with a smaller number of topics (K) than the optimal number will lead us to discover more clusters of topics (N). As we increase K, we expect N to stay relatively stable until higher Ks result in fewer N. We therefore select our parameter K to be when K=N. Alternatively, we can also calculate the perplexity. A lower perplexity score is an indication of a better model. We observe that perplexity tends to decrease rapidly before flattens out. In this approach, we choose the smallest value of K that is able to explain the data: i.e. the value of K at the point in which the perplexity flattens out (Fig. S1A).

**Fig. 1.**
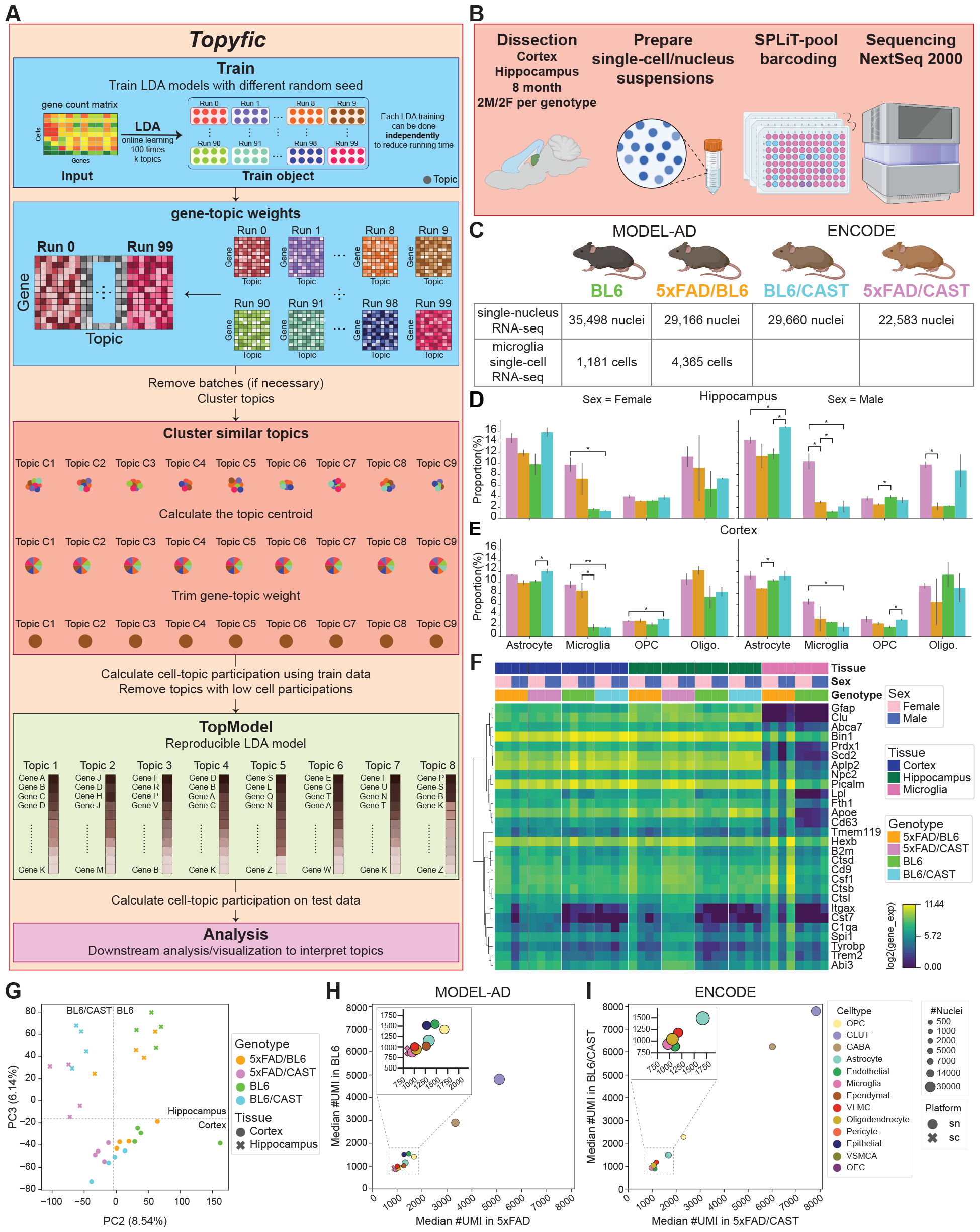
Overview of Topyfic and datasets. **A**. Overview of Topyfic workflow, as described in the text. **B**. Diagram of experimental design of single cell and single nucleus RNA-seq using Split-seq. **C**. Number of recovered nuclei/cells from each genotype after filtering. **D-E**. Proportion of glial cells recovered from a single nucleus dataset in each genotype in **D**. the hippocampus and **E**. cortex. **F**. Hierarchical clustering of gene expression markers for microglia, astrocytes, and Alzheimer’s disease (AD) marker genes in each pseudobulked sample. **G**. PCA plot of pseudobulked samples. **H-I**. Comparison of the median of UMI counts in cell types in AD mice vs. WT H. in MODEL-AD and **I**. in ENCODE datasets. Dot size reflects the number of nuclei in each cell type.

We evaluated the sensitivity of the resulting topics to important parameters. First, we investigated the effect of the number of cells on the number of reproducible topics. As expected, increasing the number of cells led to the identification of rarer and diverse gene expression programs with more topics (Fig. S1B). In the final steps of building our TopModels, we filter topics with low cell-topic participation. Increasing the minimum cell-topic participation threshold enabled us to retain topics with stronger signals in each cell (Fig. S1C). As a default criterion, we focused on topics that represent at least 1% of the gene expression in cells. We then use Leiden clustering to form our consensus topics, therefore all the inputs related to clustering, such as the resolution, which is a value controlling the coarseness of the clustering, can also be altered. Increasing this value can result in more topics, however, the defaults were found to be adequate for our analyses (Fig. S1D).

### Comparing a mouse model of AD across two genetic backgrounds

We first analyzed the overall characteristics of two related mouse brain datasets for the 5xFAD mouse model of AD(15) and matching controls for 8 month old mice before applying Topyfic. The genetic background of the 5xFAD mouse model of AD is C57BL/6J (“BL6”) and it is normally studied as a hemizygote, i.e. with only one copy of the transgene on an otherwise regular BL6 background. The first dataset consists of single-nucleus RNAseq from the cortex and hippocampus of 2 male and 2 female 5xFAD x CAST/EiJ F1 hybrid mice and matching BL6 x CAST/EiJ controls from the ENCODE consortium, while the second dataset consists of 2 male and 2 female cortex and hippocampus snRNA-seq as well as microglia single-cell RNA-seq of 5xFAD hemizygotes and matching BL6 controls from the MODEL-AD consortium (Fig. 1B-C). As all mice from ENCODE have the BL6 x CAST/EiJ F1 background, and all MODEL-AD mice have the BL6 background, we will use the consortium names and genotypes interchangeably.

All experiments were performed using the Parse Biosciences split-pool method(16, 17) which we refer to as Split-seq. Separate Split-seq experiments were performed for the ENCODE and MODEL-AD mice and were deposited in their respective online repositories. For each dataset, demultiplexing and alignment were carried out using Parse Biosciences’ split-pipe software and STARSolo(18). Scrublet(19) was employed to identify doublets in each dataset, followed by quality control (QC) using Seurat(20) (Methods). The filtering successfully recovered a combined total of 110,907 nuclei and 5,546 microglia cells (Fig. 1C), which were annotated using marker genes and label transfer with external reference data from the Allen Brain Institute dataset(21) (Fig. S2,S3 and Methods).

Glial cells such as microglia, astrocytes, and oligodendrocytes constitute a substantial fraction of the mammalian brain, representing 27.5% of the nuclei in our snRNA-seq datasets. The proportion of glial cells is influenced by several factors, including genotype, sex, and brain region. We examined the variation of glial cells by genotype and sex in each tissue separately. As expected, we found a higher portion of microglial cells in 5xFAD mice regardless of genetic background. Despite recovering more nuclei from BL6 mice, we observed a higher proportion of glial nuclei in the BL6/CAST genotype, highlighting how genetic diversity contributes to substantial differences in glial cell abundance. Interestingly, there is more variation between sexes in mice with the BL6 background compared to mice with the BL6/CAST background. In particular, 5xFAD/CAST males have similar numbers of microglia in the hippocampus when compared to 5xFAD/CAST females, which is substantially higher than 5xFAD males (Fig. 1D-E).

Expression levels of marker genes for disease-associated microglia (DAM), astrocytes, and oligodendrocytes in pseudo bulk for each mouse showed higher expression in 5xFAD versus WT and more uniformity between replicates and sexes in 5xFAD/CAST than 5xFAD in both hippocampus and cortex (Fig. 1F). Principal component analysis (PCA) confirmed that genotype contributes to major transcriptomic differences across the dataset, with PC2 (8.54%) corresponding to genotype and PC3 (6.14%) corresponding to brain region (Fig. 1G and Methods). Comparison of the median number of UMIs across cells in each cell type in AD and WT samples reveals reproducible patterns across both genotypes, such as neurons generally having more UMIs compared to glial cell types (Fig. 1H-I). Interestingly, we do not detect differences in the number of UMI between microglia whether using nuclei or whole cells (Fig. 1H).

In general, we observe a higher number of UMIs per nuclei in BL6/CAST genotype even though both consortia used similar sequencing depths. These results indicate that the differences are more likely associated with the genetic identity of mice, such as genotype, rather than technical procedures such as sequencing depth. In summary, 5xFAD/CAST males at 8 months show a higher proportion of DAM microglia that matches their female counterparts, unlike regular 5xFAD males at 8 months, which have lower proportion of DAM microglia than their female counterparts.

### Identifying topics related to cell type and cell state

We trained Topyfic using (a) 1 male replicate and 1 female replicate of WT mice from both genotypes (4 mice) and (b) 1 male replicate and 1 female replicate of 5xFAD transgene-carrying mice from both genotypes (4 mice) separately using all genes (Fig. 2A) with varying numbers of topics, ranging from K = 5 to 50. This iterative process allowed us to evaluate different K values and identify the final number of topics (N) that best captured the underlying structure in our data, which was found to be K=15 (Fig. 2B, methods). The TopModels for each genotype were aggregated to form our final TopModel with 28 topics that passed our low participation filter on the second replicates. For comparison with subsequent topics derived from different gene sets and cell types, we label these topics as asn1 through 28, where ‘asn’ stands for ‘all genes, single-nucleus’.

**Fig. 2.**
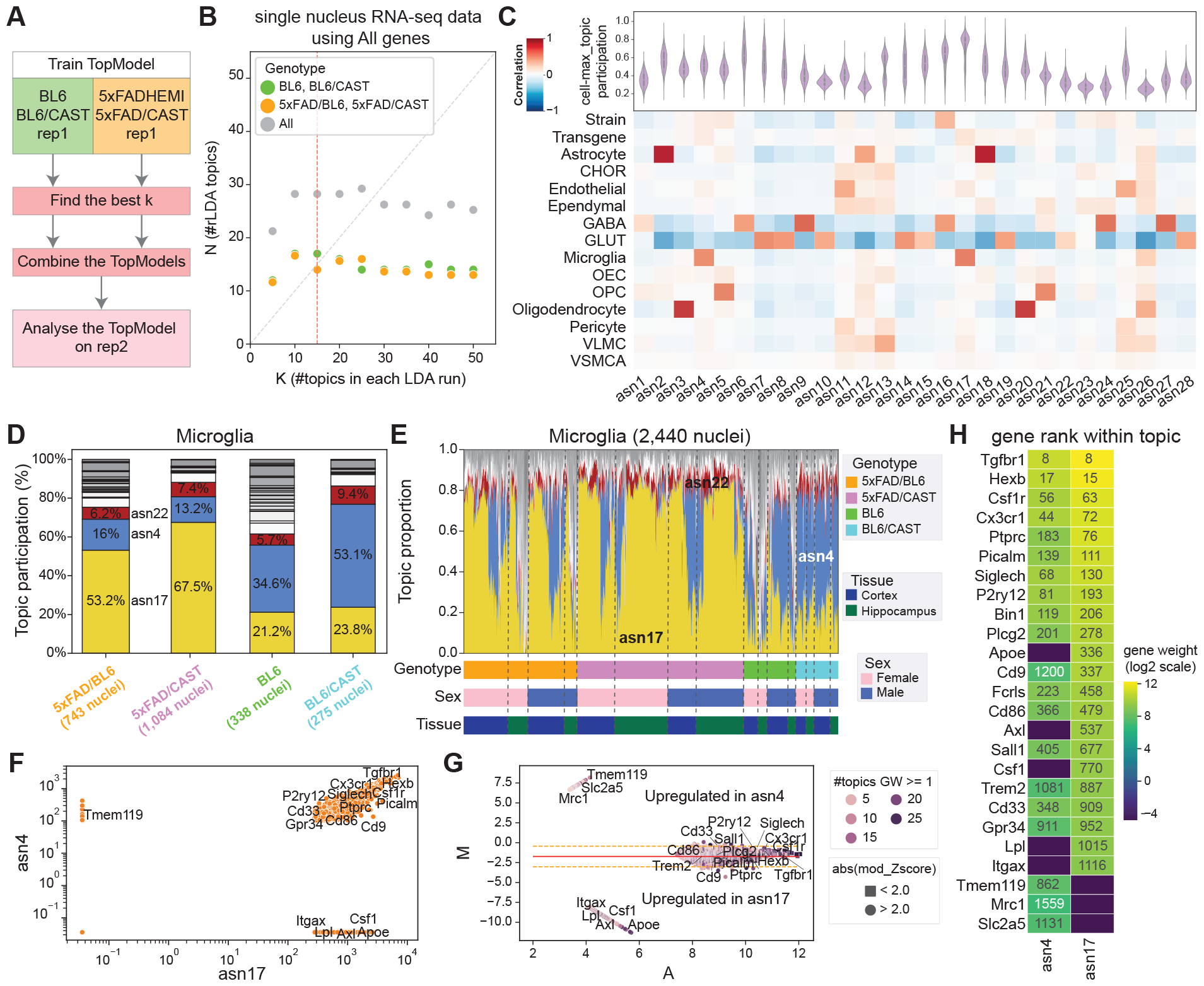
Topic modeling in single nuclei from 5xFAD/BL6 and 5xFAD/CAST cortex and hippocampus. **A**. Topics were called in 5xFAD transgenic mice and control mice separately using the first biological replicate for each mouse pair. After finding the best k to describe each training set separately, resulting topics were combined into a single TopModel, which was applied to the second technical replicate. **B**. The number of starting topics (K) versus the number of final topics (N) on the separate runs, and the combined models. Choosing K=15 for the individual runs led to a final set of 28 combined topics after filtering. **C**. Topic-trait relationship of the Spearman correlation between traits such as major cell type, strain, and transgene, across all topics. “Strain” indicates the background of mice, either BL6 (red) or BL6/CAST (blue). “Transgene” shows if the mice have the 5xFAD transgene (red) or not (blue). The violin plots show the distribution of cell topic participation whenever the topic is top ranked topic in a cell. **D**. Overall topic participation in microglia nuclei by genotype. Three major microglia topics were annotated. **E**. Structure plot of topic participation for each nucleus sorted by hierarchical clustering in each group (same genotype, sex, and tissue). **F**. Gene weights of the main two microglia topics asn4 and asn17 in log-log plot. **G**. MA plot comparing asn4 and asn17, where the X-axis (A) represents the average weight of the gene between both topics in the comparison, and Y-axis (M) represents log base 2 of the fold change of gene weight between topics. The color of each dot shows the number of topics (out of 28) where each gene has a weight above one. Modified z-score also indicated genes that were significantly differentially weighted between both topics in the comparison. **H**. Gene weights and the rank of each gene within the topic shown for asn4 and asn17. Color represents the weights of genes in the log2 scale.

We assessed the distribution of cell-topic participation, focusing on whether a topic was the predominant topic in a cell. We also performed a topic-trait relationships analysis to capture correlations between each topic and major cell types, cell states, genotype (BL6, BL6/CAST), and transgene presence (5xFAD and WT) (Fig. 2C). Topics exhibiting high cell participation consistently showed enrichment for specific cell types. Conversely, topics with low participation were generally not associated with any particular cell type. We identified topics asn4, asn17, and asn26 as corresponding to microglia, each displaying varying levels of cell-topic participation. Notably, asn17 exhibited the highest participation, while asn26 showed the lowest (Fig. 2C). Our snRNA-seq dataset includes 2,440 microglia, 77% of which are from mice with the 5xFAD transgene. The structure plot of microglia nuclei displays cell-topic participation as a stacked bar plot for each nucleus, grouped by genotype, sex, and tissue (Fig. 2D, methods). While both asn4 and asn17 are correlated with presence of the transgene, asn programs in microglia show that asn17 dominates in the transgenic mice. In mice without the transgene, microglia have a predominant mixture of cellular programs consisting of 50% asn4, 20% asn17, 7% asn26, and 23% from the remaining topics. By contrast, transgenic mice show two distinct cellular program patterns, suggesting the presence of two distinct cellular states in the mice. A minority of cells show program combinations that resemble the WT samples, which indicate a population of homeostatic microglia. The majority of microglia in transgenic mice show a significantly higher ( 65%) participation of asn17, which represents the heightened activation state of these microglia (Fig. 2E). Thus, Topyfic recovers topics representing different cell states and cell types for minor cell types such as microglia across both genotypes.

The importance of a gene’s expression to a given topic is called the gene weight. To gain insights into the differences between the two major microglia topics, we compared the weights of genes in asn4 and asn17 that have weights greater than 1. We found 120 genes specific to asn4 and 585 genes specific to asn17, as well as 1,659 genes shared between the two topics (Fig. 2F). Key genes associated with homeostatic microglia, such as *Tmem119*, exhibited significantly higher weights in asn4 compared to asn17. In contrast, genes linked to disease-associated microglia (DAM) such as *Csf1r, Itgax*, and *Apoe*, were exclusively represented in asn17 (Fig. 2F). By using an MA plot to compare topics, we found a total of 138 genes with differential weights (modified z-score > 2) in asn4, including 120 genes with large absolute log ratios (M) value (> 6.5). Conversely, asn17 displayed differential weights in 835 genes (modified z-score < −2), of which 585 had absolute log ratio (M) values > 7.5. In particular, genes overexpressed in stage 2 DAMs, such as *Apoe, Itgax, Csf1r, Lpl*, and *Axl13* were among the differentially higher weighted genes in asn17 (Fig. 2G). Genes related to microglial cell identity, such as *Tgfbr1* and *Hexb*(22), shared similar ranks in both topics, even though they had higher weights in asn17. In contrast, genes primarily expressed in homeostatic microglia such as *Tmem119* and *Slc2s5* were exclusively represented in asn4, whereas DAM genes *Apoe, Itgax*, and *Csf1r* only had significant weights in asn17 (Fig. 2H). Thus, genes with shared or specific weights in a topic can be correlated to the known underlying biology, as demonstrated in this case by the microglial neuroinflammatory signatures in mice with the 5xFAD transgene.

### Recovering topics for different activation levels in microglia scRNA-seq

Having demonstrated its performance and utility on snRNA-seq from tissues, we applied Topyfic to our complete microglia single-cell data from 5xFAD and matching BL6. After training the TopModel with multiple values of K, we selected K=5, yielding 6 topics labeled sc1sc6 (Methods). Each topic is the top participating topic in a subset of microglia (Fig. 3A). While calculating the correlation between each topic and sex did not reveal any sexspecific topics (Fig. 3B), analyzing the contribution of each topic in each genotype uncovered differences in activity between the genotypes (Fig. 3C). The structure plot illustrates three distinct cellular programs. The first combination of programs is high in sc6 with high levels of *Csf1*, and is more prevalent in 5xFAD than in BL6. The second combination of programs contains a relatively consistent proportion of sc2, sc3, and sc5 in both genotypes, suggesting a closer association with homeostatic microglia. A subset of these homeostatic cells also have participation of sc6 and low levels of sc1. The third combination of programs is more pronounced in 5xFAD mice and is primarily composed of sc1, with significantly lower participation of sc3 and sc5 compared to the previous program, indicating a stronger association with activated microglia (Fig. 3D).

**Fig. 3.**
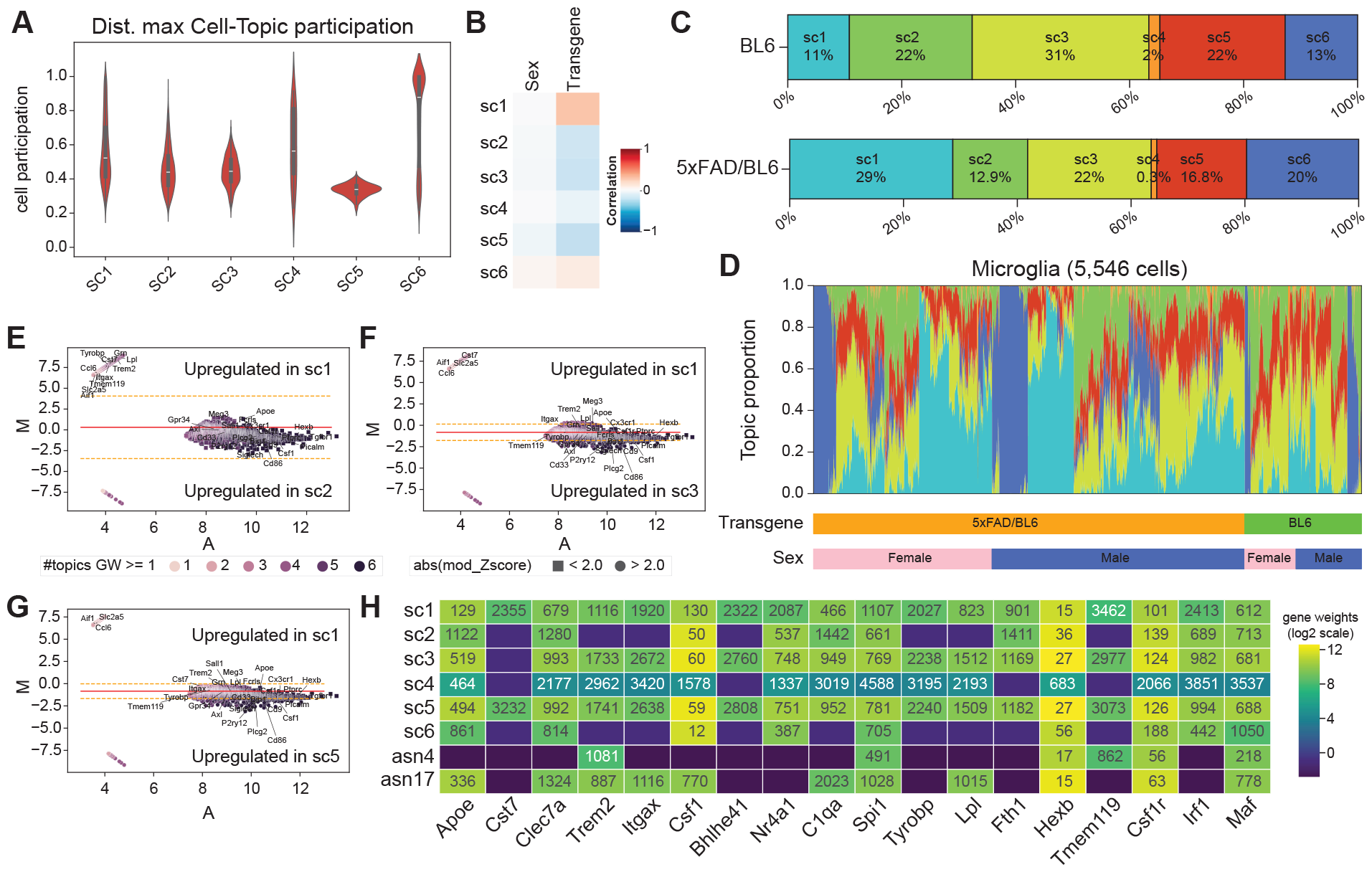
Topic modeling in scRNA-seq of microglia. **A**. Distribution of maximum cell-topic participation in each cell in each topic. B. Topic-trait correlation between sex or transgene and each topic. **C**. Topic participation broken down by genotype. **D**. Structure plot of microglia cells sorted by genotype and sex show topic participation in each cell. **E-G**. Comparison of gene weights for sc1 (activated microglia topic) **E**. versus sc2, **F**. versus sc3, and **G**. versus sc5. Color represents the weights of genes in the log2 scale. **H**. Gene weights of genes in interest in log2 scale with their rank.

Comparison of genes with weights >1 between sc1 and sc2 revealed differential weights in 1,794 genes, with only 24 genes in sc2, including the *miR-155* host gene (Fig. 3E). MicroRNAs (miRNAs) play a role in modulating inflammatory responses in microglia, and their profiles are altered in AD. Notably, the pro-inflammatory miRNA, *miR-155*, shows increased expression in the AD brain(23). We found disease-associated microglia (DAM) genes such as *Trem2, Apoe, Itgax, Clec7a, Axl*, and *Lpl*, alongside typical microglia gene markers such as *Tmem119, Olfml3*, and *Cd68* with higher weights in sc1 (Fig. 3E). A comparison between sc1 and sc3 reveals 562 genes with higher weights in sc1 (modified zscore > 2) and 247 genes with higher weights in sc3 (modified z-score < −2) (Fig. 3F). Primed microglia have the potential to induce the production of amyloid *β* (A*β*), tau pathology, neuroinflammation, and reduce the release of neurotrophic factors. This can lead to the loss of normal neurons in both quantity and function, a phenomenon strongly associated with AD. Genes such as *Cst7* and *Slc2a5*, part of the primed microglia pathway(24), were upregulated in sc1. Comparison between sc1 and sc5 reveals 465 genes with weights higher in sc1 (modified z-score > 2) and 280 genes with weights higher in sc5 (modified z-score < −2) (Fig. 3G). In all three comparisons, genes associated with disease-associated microglia are higher in sc1 compared to the three homeostatic programs. Weights and ranks of homeostatic and DAM genes across microglial cells and microglial nuclei are similar between topics sc1 and asn17. Genes such as *Lpl, Apoe*, and *Trem2* exhibit higher gene weights and lower ranks in sc1 and asn17 compared to the rest of the single-cell topics, including microglial topic asn4 (Fig. 3H). Lipoprotein lipase (*Lpl*), the rate-limiting enzyme in lipoprotein hydrolysis, is predominantly expressed in microglia such as phagocytic disease-associated microglia thought to be protective in AD(25). Increased expression of genes such as *ApoE, Trem2*, and *Lpl* in microglia during development, damage, and disease suggests that increased lipid metabolism is needed to fuel protective cellular functions such as phagocytosis(26). Thus, Topyfic recovered an activated microglial state topic using either single-cell or single-nucleus RNA-seq, with the main difference corresponding to how scRNA-seq topics capture multiple subtypes of homeostatic microglial programs.

### Topics derived from regulatory genes are sufficient to define cell types and cell states

While numerous genes are used as markers for distinct cell types and states, we hypothesized that cellular programs are fundamentally constructed from a core set of regulatory genes. Therefore, we explored identifying cellular programs using a restricted LDA vocabulary of regulatory genes. Transcription factors and other genes based on Gene Ontology (GO) term annotations were chosen based on their impact on transcriptional regulation, including known regulatory genes such as the Id family (inhibitors of DNA binding and cell differentiation, despite lacking a DNA binding domain themselves)(27) (Methods). Overall, TFs constitute approximately 50% of the 2,701 genes included in our regulatory gene list (Fig. 4A). This approach aims to elucidate impactful cellular programs using a curated vocabulary.

**Fig. 4.**
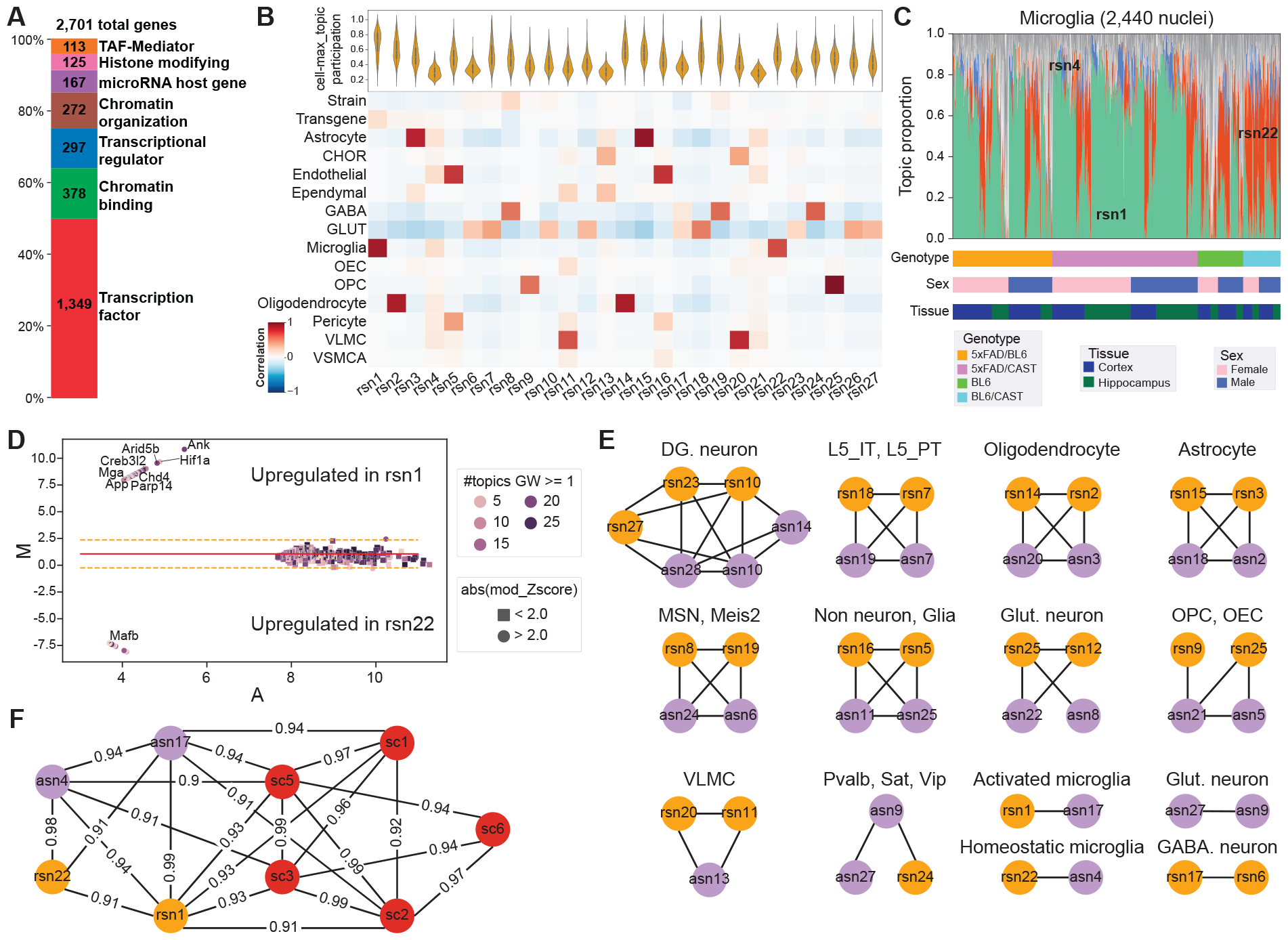
Topic modeling using regulatory genes. **A**. Breakdown of regulatory gene categories in mouse. **B**. Distribution of cell-topic participation in single-nucleus datasets. Topics rsn1 and rsn22 are enriched in microglia, with rsn1 also enriched in transgenic mice. **C**. Structure plot for microglia nuclei sorted by genotype and sex. rsn1 mostly contributes to the AD mice, whereas rsn22 is primarily found in WT mice. **D**. MA plot comparison of gene weights between rsn1 and rsn22. The color of the dots represents the number of topics with the gene weight higher than the pseudocount, and the style of each dot indicates if the difference between gene weights is significant (circle) or not (square) based on a modified z-score of M values. **E**. Topic cosine similarity > 0.9 between topic pairs using all genes and regulatory topics. **F**. Topic cosine similarity > 0.9 between microglia topics from single nucleus (asn), regulatory genes (rsn), and microglia single-cell (sc).

We trained Topyfic models on 52,685 nuclei across 2,701 regulatory genes using various K values as described previously, and once again combined models with K = 15 to obtain 27 reproducible topics labeled as rsn (regulatory single nucleus). Analysis of topic-trait relationships showed that individual topics are highly correlated to specific cell types, 5xFAD transgene presence, or genotype. For instance, both rsn1 and rsn22 correspond to microglia while rsn1 also correlated to the transgene presence, rsn3 and rsn15 to astrocytes, rsn2 and rsn14 to oligodendrocytes, and a dozen topics correspond to different neuronal subtypes (Fig. 4B). Similarly to the results using all genes, topics with the highest maximum cell participation are consistently enriched for specific cell types or cell states. By contrast, topics with low maximum cell participation are typically not associated with any particular cell type (Fig. 4B). At our chosen resolution, all cell types with > 350 nuclei (0.32%) are associated with at least one topic. In summary, our topics demonstrate a striking alignment with our annotated cell types.

The structure plot of microglia nuclei shows that as cells activate, rsn22 is gradually replaced by rsn1 while minor topics remain constant (Fig. 4C). Comparing the gene weights of the two microglia topics reveals remarkably similar topic compositions, aligning with our expectations (Fig. 4D). Only a few genes exhibit an absolute modified z-score value > 2 (93 in rsn1, 15 in rsn22) (Fig. 4D). Among the 93 genes upregulated in rsn1, microglia gene markers such as *Ank, Hif1a, Arid5b, Creb3l2*, and *Srpk2* show the highest modified zscores. The expression of serine/threonine-protein kinase 2 (*Srpk2*) is associated with the production of proinflammatory cytokines and M1 polarization of microglial cells, suggesting a potential connection to the cognitive decline observed in the AD mice model(28). *Ank* is a membrane-phosphate transporter that has a microRNA, *Mir7117*, embedded in its intron. *Ank* exhibits the highest absolute log ratio (M) value (10.8) and is also upregulated in laser-captured microglia in the brains of individuals with AD(29). In summary, regulatory genes alone can differentiate between homeostatic and disease-associated microglial states.

We evaluated the similarity between topics learned using all genes (asn) and regulatory genes (rsn) using cosine similarity (methods). Using a similarity threshold of > 0.9, we identified 14 clusters of highly correlated topics that matched asn topics to rsn topics (Fig. 4E). As anticipated, topics representing common cell types were found to cluster together with at least one topic from each method type. At the selected threshold, activated microglia topic rsn1 only matches topic asn17, whereas homeostatic microglia rsn22 only matches the corresponding asn4. We further clustered the nine microglia topics (2 asn, 6 sc, 2 rsn) and found a higher correlation of activated microglia topic sc1 with asn17 and rsn1 than with the other asn and rsn topics (Fig. 4F). Thus, the expression patterns of regulatory genes alone are adequate to define cell types in the brain and states of microglial activation.

## Discussion

We developed a grade of membership (GoM) model using Latent Dirichlet allocation (LDA) called Topyfic and applied it to mouse single-cell and single-nucleus RNA-seq brain datasets to infer topics that capture cell types, subtypes, and cell states. Current implementations of LDA for analyzing single-cell RNA-seq data do not consider the stability and consistency of the model(30–33). A robust LDA model should be less sensitive to variation in the initial conditions, such as different random seeds. To achieve this, we implemented reproducible LDA (rLDA) in Topyfic, which automatically runs LDA multiple times and aggregates similar topics. In particular, our strategy identifies the optimal number of topics at the given resolution. We then score topic participation in the whole dataset, which enables us to anchor each topic to a specific cell type, guiding the subsequent inference of the global topic distributions over genes to prioritize genes differentially weighted in each topic.

We applied Topyfic to mouse brain tissues and microglia cells in control and well established and thoroughly examined AD mouse models such as 5xFAD to validate our strategy. We recovered topics that reflect different cell types, including glia with different levels of activation. Interestingly, our findings indicate increased DAM microglia in male 5xFAD/CAST F1s compared to male 5xFAD/BL6 mice. While we recover the expected sex-specific difference in the higher amount of detected DAM microglia in female 5xFAD/BL6 mice compared to male 5xFAD/BL6 mice as previously reported(15), this pattern is not seen in 5xFAD/CAST F1s, where both sexes show an equal proportion of DAM microglia. This suggests that genetic variation affects disease progression in the different sexes in mouse models of AD. While previous studies have compared APP/PS1 transgene expression in inbred wild-derived strains such as CAST also at 8 months of age and found similar levels of microglia activation in transgenic CAST and transgenic BL6, the study only analyzed females(34). Whether it is 5xFAD/CAST F1 males that are atypical because of their increased activation or alternatively 5xFAD/BL6 inbred males that are atypical because of their lower activation remains to be determined.

We have shown that Topyfic recovers topics related to cell types or activities, including multiple topics relating to distinct activation states in microglia. We also show that using regulatory genes is enough to identify cell types or cell states by limiting our dictionary of genes to those that are the most likely to be directly involved in transcriptional and posttranscriptional regulation. For example, the top differential gene in topic rsn1 was *Ank1*, which has a 4-fold upregulation in AD microglia(29). While *Ank1* is a structural protein, it is also the host gene of *Mir7117*, which suggests that the microRNA could be playing a role in AD. This potential role for *Ank1* as a microRNA host gene in topic-based cellular programs would have been missed in traditional protein-coding gene marker analysis.

Cellular programs defined using topic modeling have two major benefits over traditional cluster-based approaches to single-cell analysis. First, each gene can contribute more than one topic with different weights, which is more reflective of the pleiotropic nature of gene activity. The second substantial benefit is that each nucleus can have a unique linear combination of topic membership, rather than force it to be only part of a homogeneous cluster(20, 35). This approach infers a higher and more abstract level of transcriptional activity, represented as topics. The latent topic structure is biologically interpretable, as cells carry out their functions by engaging simultaneously in multiple cellular programs related to cell identity, activation state, cell cycle, or circadian rhythm. Each pathway relies on different amounts of a gene’s product, and the overall gene expression reflects the combined requirements across all topics. This GoM representation of cell-topic participation can be interpreted as a dimensionally-reduced portrayal of a cell’s distinct transcriptional activities, which is what we are truly interested in elucidating when applying single-cell techniques to study already well-surveyed samples that have been previously characterized by bulk techniques. The sooner the field moves from individual gene marker-based analyses to cellular-program based approaches, the more likely we are to obtain useful biological insights to match our wants and needs.

## ACKNOWLEDGEMENTS

A.M., B.J.W., L.R were supported by UM1HG009443 (ENCODE). A.M. was also supported by NIA U54 AG054349 (MODEL-AD).

## AUTHOR CONTRIBUTIONS

A.M. and N.R. designed the package and study and wrote the manuscript. N.R. wrote Topyfic and its documentation, performed data analysis, and generated figures. E.R. performed Parse Biosciences snRNA-seq experiments for the ENCODE samples and preprocessed the data. B.A.W. processed the tissues for the ENCODE samples. H.Y.L. performed Parse Biosciences sc and snRNA-seq experiments for the MODEL-AD samples. All co-authors edited the manuscript.

## DATA AVAILABILITY

- Mouse snRNA-seq dataset from ENCODE
- Mouse sn/scRNA-seq dataset from MODEL-AD

## CODE AVAILABILITY

- Topyfic package
- Data processing and figure generation code

## Materials

### Mice and tissue collection

All mice were housed following the guidelines outlined in the Guide for Care and Use of Laboratory Animals. Approval for all experimental procedures was obtained from UCI’s Institutional Animal Care and Use Committee (IACUC), adhering to both institutional and national guidelines. Model-AD samples were obtained from 5xFAD/BL6 mice (Tg(APPSwFlLon,PSEN1*M146L*L286V)6799Vas/Mmjax, RRID: MMRRC_034840-JAX) covered under the IACUC protocol #AUP-21-100 and bred by the Transgenic Mouse Facility at UCI. Left cortex and left hippocampus tissues from 8 month old mice were snap frozen in liquid nitrogen at UCI and stored at −80°C. ENCODE samples were obtained from 5xFAD x CAST/EiJ (RRID: IMSR_JAX:000928) F1 hybrids covered by IACUC protocol #IA21-1647 and bred by Jackson Laboratories (JAX). Left cortex and left hippocampus tissues from 8 month old mice were snap frozen in liquid nitrogen at JAX and shipped to UCI on dry ice.

### Single-nucleus isolation and fixation

All single-nucleus samples regardless of genotype or tissue were processed identically. On ice, tissues from each mouse were transferred to a chilled gentleMACS C Tube (Miltenyi Biotec cat. #130-093-237) with 2 mL Nuclei Extraction Buffer (Miltenyi Biotec cat. #130-128-024) supplemented with 0.2 U/µL RNase Inhibitor (NEB cat. #M0314L). A gentleMACS Octo Dissociator (Miltenyi Biotec cat. #130-095-937) was used to dissociate nuclei from whole tissues. The resulting suspensions underwent rounds of filtering through mesh strainers (70 µm, Miltenyi Biotec cat. #130-110-916, then 30 µm, #130-098-458). Finally, nuclei were resuspended in PBS + 7.5% BSA (Life Technologies cat. #15260037) and 0.2 U/µl RNase inhibitor and kept on ice. Manual counting was performed using a hemocytometer and DAPI stain (Thermo cat. #R37606). After counting, nuclei were fixed using Parse Biosciences’ Nuclei Fixation Kit v1 (Parse Biosciences cat. #WN100), following the manufacturer’s protocol. Between 1 and 4 million nuclei per sample were incubated in fixation solution for 10 minutes on ice, followed by permeabilization for 3 minutes on ice. The reaction was quenched and nuclei were centrifuged and resuspended in 300 µL Nuclei Buffer (Parse Biosciences cat. #WN101) and DMSO (Parse Biosciences cat. #WN105). The fixed samples were assessed under a microscope and manually counted as previously described. Aliquots of fixed nuclei were slow-frozen in a Mr. Frosty (Thermo cat. #5100-0001) and stored at −80°C.

### Microglia single-cell isolation and fixation

Freshly prepared tissues were used for microglia isolation. Perfused right cortex and hippocampus were dissociated together using the Adult Brain dissociation kit (Miltenyi Biotec cat. #130-107-677) and gentleMACS Octo Dissociator (Miltenyi Biotec cat. #130-095-937) with heating. The resulting suspension was filtered with a 70µm mesh strainer (Miltenyi Biotec cat. #130-110-916). Debris were removed using the debris removal solution from the dissociation kit. Myelin were removed from the single-cell suspensions using negative selection with Myelin Removal Beads II (Miltenyi Biotec cat. #130-096-733) and LS columns (Miltenyi Biotec cat. #130-042-401). The resulting cells were enriched for microglia with magnetic labeling and positive selection using CD11b MicroBeads (Miltenyi Biotec cat. #130-093-634) and LS columns. Isolated microglia eluted in 1.8mL of bead buffer (0.5% BSA in DPBS) from the LS columns were centrifuged at 550 xg for 10 min at 4°C. Cell pellet with 10-15uL bead buffer was resuspended in 730µL of Cell buffer (Parse Biosciences cat. #WF101) from the Parse Biosciences kit (V1.3.0) with 0.5% BSA (Life Technologies cat. #15260037), and counted (1:4 dilution) with TC20 Automated Cell Counter (Bio-rad cat. #1450102). Cells were fixed using Parse Biosciences’ Nuclei Fixation Kit v1 (Parse Biosciences cat. #WF102), following the manufacturer’s protocol. Between 150000 - 800000 cells per sample were incubated in fixation solution for 10 minutes on ice, followed by permeabilization for 3 minutes on ice. The reaction was quenched and nuclei were centrifuged and resuspended in 150 µL Cell Buffer. The fixed samples were counted using TC20 automated cell counter. DMSO (Parse Biosciences cat. #WF105) was added to the fixed cells and cells were slow-frozen in a Mr. Frosty (Thermo cat. #5100-0001) and stored at −80°C.

## Methods

### Parse Biosciences Split-seq Experiments

The Model-AD and ENCODE libraries were prepared in two separate experiments using Parse Biosciences’ Evercode WT Kit v1 (cat. #EC-W01030), one kit per experiment, following the manufacturer’s protocol. Fixed samples were thawed and added to the Round 1 barcoding plate at 15,000 nuclei/cells per well when possible across 48 wells. For microglia samples with low numbers, the entire sample was added. Each tissue sample from one individual was loaded into a single well. RNA was reverse transcribed in the fixed nuclei/cells using oligodT and random hexamer primers and the first barcode was annealed. After RT, samples were pooled and randomly distributed across 96 wells of the Round 2 ligation barcoding plate for in situ barcode ligation. After Round 2, samples were pooled and randomly redistributed into 96 wells of the Round 3 ligation barcoding plate for ligation of the third cell barcode and Illumina adapters. Finally, samples were counted with a hemocytometer and distributed into 6 subpools of 15,000 nuclei for a target of around 75,000 nominal nuclei/cells per tissue. To prepare libraries, the nuclei/cells in each subpool were lysed and the barcoded cDNA was amplified. The cDNA was purified with AMPure XP beads (Beckman Coulter cat. #A63881) and quality checked with the Qubit dsDNA HS Assay Kit (Thermo cat. #Q32854) and Bioanalyzer 2100 (Agilent cat. #G2939A) High Sensitivity DNA Kit (Agilent cat. #5067-4626). Subpool cDNA (100 ng) was fragmented and Illumina P5/P7 adapters were ligated during the last amplification, followed by size selection and quality check with the Bioanalyzer and Qubit. An Illumina NextSeq 2000 and P3 200 cycles kits (Illumina cat. #20040560) were used to sequence libraries with 5% PhiX spike-in with as paired-end, single-index reads (115/86/6/0) to an average depth of 187 M reads per Model-AD library, and 183 M reads per library.

### Datasets

One microglia single-cell RNA-seq dataset of cortex and hippocampus at 8 months on 5xFAD mouse model36 and matching wild type (C57BL/6J) were used to demonstrate Topyfic behavior on microglia cells. We also combined two single-nucleus RNA-seq datasets of cortex and hippocampus from the 5xFAD mouse model of AD in two different genetic backgrounds (B6J and B6CASTF1/J) from the Model-AD and ENCODE consortiums, respectively. The Model-AD snRNA-seq was performed in 8 month old 5xFAD and matching wild type (C57BL/6J) mice, and ENCODE snRNA-seq was performed in 8 month old 5xFAD/CAST and matching WT (B6CASTF1/J) hybrid mice.

### Preprocessing scRNA-seq and snRNA-seq data

Raw fastq files were processed using Parse Bioscience’s splitpipe software (v1.0.3p) to assign reads to single cells and nuclei. In order to provide sample-level fastqs to the ENCODE portal, all data was demultiplexed using the sample-level barcode (barcode 1) from the output of split-pipe to be aligned and quantified with the ENCODE uniform processing pipeline. We use STARSolo with GeneFull_Ex50pAS settings and the GENCODE vM21 annotation to generate UMI count matrices, annotated using GENCODE vM21. We removed low-quality cells using a UMI cutoff of 500 based on the knee plots and then performed Scrublet(19) doublet detection. Cells and nuclei with <500 or >30,000 UMIs, more than 500 genes, and a doublet score >0.2 were removed in downstream analysis. In addition, nuclei were required to have a mitochondrial gene expression score of <0.5%, while cells had a more lenient threshold of <5%. Seurat17 V4 was used to perform normalization, UMAP dimensionality reduction, and clustering. Each dataset (Model-AD 5xFAD and B6J WT snRNA-seq, ENCODE 5xFAD/CAST snRNA-seq, and Model-AD scRNA-seq microglia) were preprocessed and clustered separately, with 50, 40, and 15 clusters, respectively, after removal of 2 doublet-high clusters from the Model-AD snRNA-seq dataset and 3 doublet-high clusters from the ENCODE dataset(Fig. S2,S3).

To facilitate cell type annotation, a downsampled version of the 1M whole cortex and hippocampus 10x atlas from 8 week old mice available on the Allen data portal(17) was used to transfer subtype-level annotations using “FindTransferAnchors” in Seurat. Each cluster was then manually annotated using the resulting Allen Atlas labels and marker gene expression. Overall, this process identified 13 major cell types and 34 subtypes in the snRNA-seq data, and 5 cell types (75% of which are microglia) in the scRNA-seq data. After defining annotations for each dataset, we extracted the raw gene count matrices along with gene and sample information from Seurat objects and embedded all information into the Anndata(36) file format where gene information is ‘var’ and cell information is ‘obs.’ Finally, we applied depth normalization(37) individually to each dataset to prepare them for the rest of the analysis (Fig. 1A).

### Selection of regulatory genes

Regulatory genes were determined by microRNA-host gene correlations, annotated transcription factors, and genes annotated with Gene Ontology (GO) terms based on their impact on transcriptional and chromatin regulation. GO term-derived genes were collapsed into 5 categories: histone modifiers, from GO terms related to histone acetyltransferases (GO: 0004402), histone deacetylases (GO: 0004407), histone methyltransferases (GO: 0042054), and histone demethylases (GO: 0032452); TBP-associated factors and members of the Mediator complex (TAF-MED, GO: 0016592 and GO: 0006352), chromatin binding (GO: 0003682), chromatin organizers (GO: 0006325 and GO: 0030527), and transcription regulators (GO: 0140110). The final list of 2,701 expressed regulatory genes has 7 biotype categories including microRNA host genes and transcription factors.

### Input data to Topyfic

Topyfic accepts input in the form of a preprocessed expression matrix embedded within the Anndata format(36). This format contains gene information as ‘var’ and cell information as ‘obs.’ Users can generate this input format from the output of popular single-cell tools like Scanpy(35) or Seurat(20). For topics modeling using the regulatory gene set, the expression matrix was subset for the genes of interest, then normalized and formatted. It’s important to note that Topyfic leaves the choice of performing normalizations to the user’s discretion. However, it is strongly recommended, especially when dealing with data originating from different technologies. Without normalization, there is a heightened risk of detecting topics influenced by batch differences, even when they don’t represent meaningful biological signals. To mitigate this issue, we apply depth normalization37 individually to each dataset. This approach effectively implements depth normalization and variance stabilization, enabling the accurate identification of recurring patterns while minimizing the impact of technical variations.

### Topic modeling using Latent Dirichlet Allocation (LDA)

Topic modeling is a type of statistical model that uses unsupervised machine learning to identify groups of similar words in each document. LDA is a generative probabilistic Bayesian model that operates on the assumption that documents can be represented as random mixtures over latent topics, where each topic is characterized by a distribution over a set word vocabulary. In simpler terms, each document is a mixture of topics, and each topic is a mixture of words, where words can be repeated in different topics with different weights. In the context of single cell/nucleus RNA-seq data, cells correspond to documents, genes to words, and counts are equivalent to word frequencies. We hypothesize that there are recurring latent patterns or “topics” in count data such as large gene expression matrices. Topics are composed of genes with distinct weights that can together recreate underlying patterns of gene expression profiles for each individual cell.

### LDA Model Training

We employed scikit-learn’s LDA implementation (v1.3)(38) with options to allow users to change default parameters including batch size and learning method based on their data (default: learning_method=online variational Bayes method, batch_size=1000, max_iter=10). Due to the random initialization of LDA algorithms, topic definitions can vary substantially each time that the algorithm is rerun, which hinders their interpretability. Therefore, we train the LDA model with several distinct random seeds (default 100 times) to capture all possible topics. After training all LDA models, we built our gene-topic weight matrix using all the obtained models. Even though learning each LDA model is not overly time-consuming, learning it 100 times can be time-intensive and will increase by increasing the number of input data(cells/nuclei). To reduce run time, we have added another feature to train each LDA model separately and then combine all of them to make the final LDA train object.

### TopModel Construction

We assume topics that are independent of the random seeds should have a similar gene weight profile. Leveraging this hypothesis, we employ the Leiden algorithm to cluster all topics with similar gene weight profiles. In cases where batch effects may be present, we incorporate Harmony(39) for batch correction, ensuring that the topics remain consistent across different datasets. Once the clusters are defined, Topyfic calculates the topic centroid (mean of gene weights) for each cluster to create a new gene weight matrix. Then it will trim the matrix by calculating 90% of the cumulative sum of gene weight and reassign the rest to a pseudocount (1 divided by the total number of topics), creating a new reproducible LDA model (TopModel). To enhance the quality of topics, we implemented a filtering step that discards small clusters of topics based on cell-topic participation, with the default threshold set at less than 1% of cells. If any topic is eliminated in this step, Topyfic will redo the trimming and reassign the rest to the new pseudocount. This filtering step aids in eliminating topics that may have emerged due to random seed fluctuations, focusing our analysis on the more stable and biologically meaningful topics.

### Annotating topics

A topic is essentially a vector of gene weights denoting their relative contribution to that specific topic. It is important to note that a single gene can appear in multiple topics with different weights. Topyfic offers different functional enrichment analyses for each topic, enhancing its utility and our understanding of each topic. These analyses encompass Gene Ontology (GO) analysis, Gene Set Enrichment Analysis (GSEA)(40), and pathway analysis using the REACTOME database(41). Furthermore, Topyfic supports the comparison of the two topics. This is achieved by transforming the data onto two scales: M (log ratio) and A (mean average). These scales facilitate the calculation of a modified z-score based on the M value, allowing for meaningful comparisons between topics in terms of their gene weights.

### Analysis object

The Analysis object is a pivotal component in the post-processing phase of Topyfic, aiding in biological interpretation of the topics and data following training of the TopModel. After successfully training the TopModel, analysis of the TopModel itself commences. This analysis includes the calculation of ‘cell-topic participation,’ which quantifies the extent to which each topic contributes to each cell. In essence, it represents probability distributions for each row, ensuring that the sum of topic participation for each cell equals one. To facilitate a comprehensive understanding of the topics and their relationships, Topyfic offers several visualization tools.

Using grade-of-membership models helps us estimate membership proportions for each cell/nucleus in each topic, visualized as a structure plot(42, 43). The structure plot displays the estimated membership proportions of each cell/nucleus as a stacked bar plot, with different colors representing different reproducible topics. To enhance the visualization of inferred cellular programs from the data, cells are sorted within selected traits in a given order. Within each trait group, cells are further ordered based on their similarity in estimated membership proportions, employing Ward’s linkage(44). A pie chart summarizes the structure plot, providing a representation of the overall contribution of each topic of all the cells/nuclei in a trait group. A topic-trait relationship heatmap visualizes the Spearman correlation coefficients between each trait and topic as a heatmap. These visualizations serve as valuable aids in interpreting the topics and their associations with other relevant traits or characteristics.

### Comparing Topics

An advantage of employing LDA to discover topics lies in the representation of gene weights as probability distributions within each topic, ensuring that the sum of gene weights in any topic equals one. Leveraging this property, Topyfic offers a valuable feature for comparing topics based on their gene membership profiles. To facilitate this comparison, Topyfic normalizes the gene weights and assesses the similarity between any pair of topics in the gene membership space. This similarity evaluation can be performed using various methods, including Pearson correlation, Spearman correlation, cosine similarity, and information-theoretic metrics like the Jensen–Shannon divergence. Once these comparisons are completed, the results can be visualized as a graph or heatmap through Topyfic. This visualization allows for a clearer understanding of the relationships between different topics based on their gene memberships.

### LDA parameter settings

In addition to determining the final number of topics, other parameters may require tuning based on the input data. We suggest using ‘online’ as a learning method, which uses an online variational Bayes method. This method updates the gene weight in each topic during each EM update using a mini-batch of training data. We can also tune learning_decay which controls learning rate in the online learning method. Besides these, other possible search parameters could be batch_size (number of cells to use in each EM iteration; default=1000) and max_iter (the maximum number of passes over the training data, aka epochs; default=10). Given enough computing resources, it might be worthwhile to experiment with these parameters.

### Pseudobulk calculation

A pseudobulk sample is formed by aggregating the expression values that pass QC from groups of nuclei originating from the same individual, which represents the experimental unit of replication.

### Principal component analysis (PCA)

Principal component analysis was performed through scikit-learn on the pseudobulk matrix, where 27 components explained 95% of the variance in the single nucleus data.

### Topyfic analysis of single-nucleus RNA-seq data

The normalized gene-count matrix was divided based on samples with and without the 5xFAD transgene for training the TopModel through Topyfic. Initial TopModel training was performed on the first replicate of the 5xFAD and WT samples separately, employing default parameters except minimum cell participation which was 0.5% of the total number of nuclei (120.7 for the 5xFAD dataset and 142.725 for the WT dataset) using different numbers of topics (K) ranging from 5 until 50. K=15 was chosen for further analysis. Subsequently, Topyfic was used to combine TopModels and remove topics with cell participation lower than 0.5% of the total number of nuclei in both datasets (321.11) to obtain the final reproducible topics. To demonstrate that the TopModel learned meaningful topics that could be used to analyze other related datasets, we applied the trained TopModel to the second replicate of the single-nucleus data.

### Topyfic analysis of single-cell RNA-seq data

A processed gene-count matrix containing only microglia cells was passed as input to Topyfic. The TopModel was trained with all default parameters, except for the minimum cell participation which was set to 0.5% of the total number of nuclei (27.73) across different numbers of topics (K).

## Supplementary Figures

**Fig. S1.**
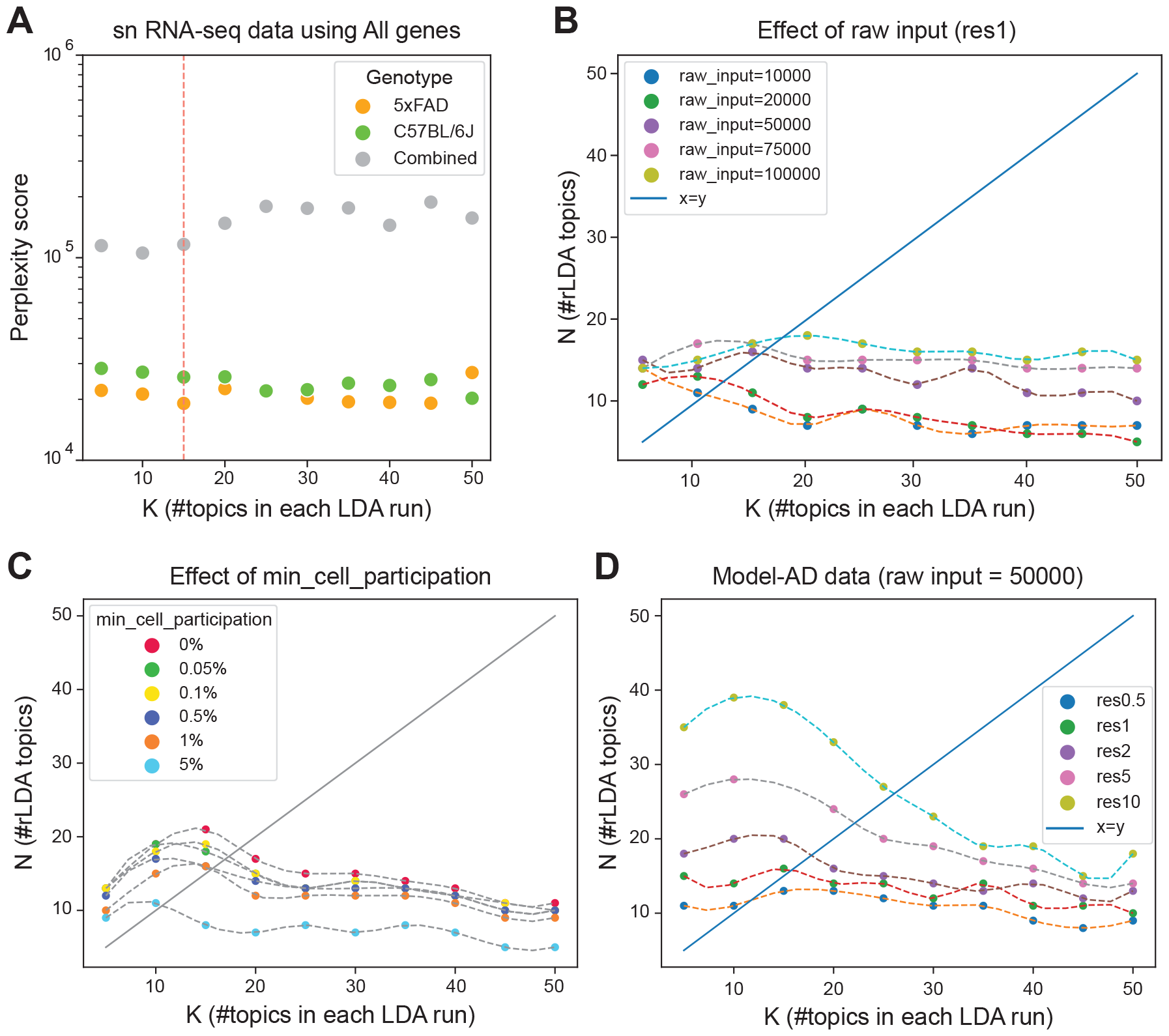
Impact of various data and Topyfic parameters on the number of topics(K). **A**. The perplexity score is based on the number of starting topics (K) on the separate runs, and the combined models. Choosing K=15 for the individual runs to have lower the perplexity score. **B**. Influence of the number of nuclei/cells on the recovered topics (K). Higher number of nuclei/cells results in a greater number of topics (K). **C**. Impact of increasing minimum cell-topic participation on the removal of smaller topics, leading to a reduction in the number of topics (K). **D**. The effect of increasing resolution on generating more clusters, subsequently increasing the number of topics (K).

**Fig. S2.**
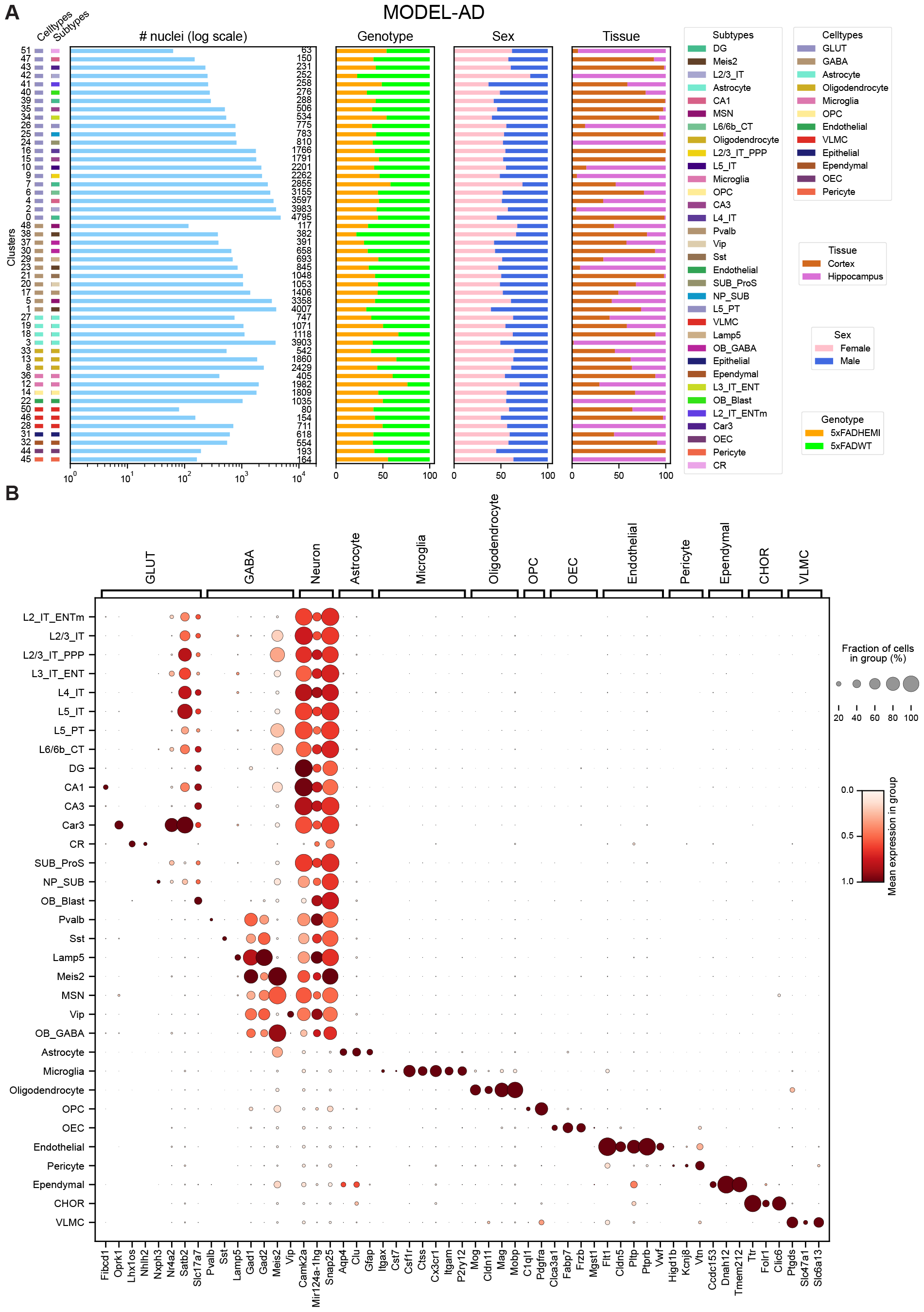
Overview of MODEL-AD dataset. **A**. Breakdown of nuclei, cell type, subtype, genotype, sex, and tissue across 52 clusters. **B**. Expression of marker genes

**Fig. S3.**
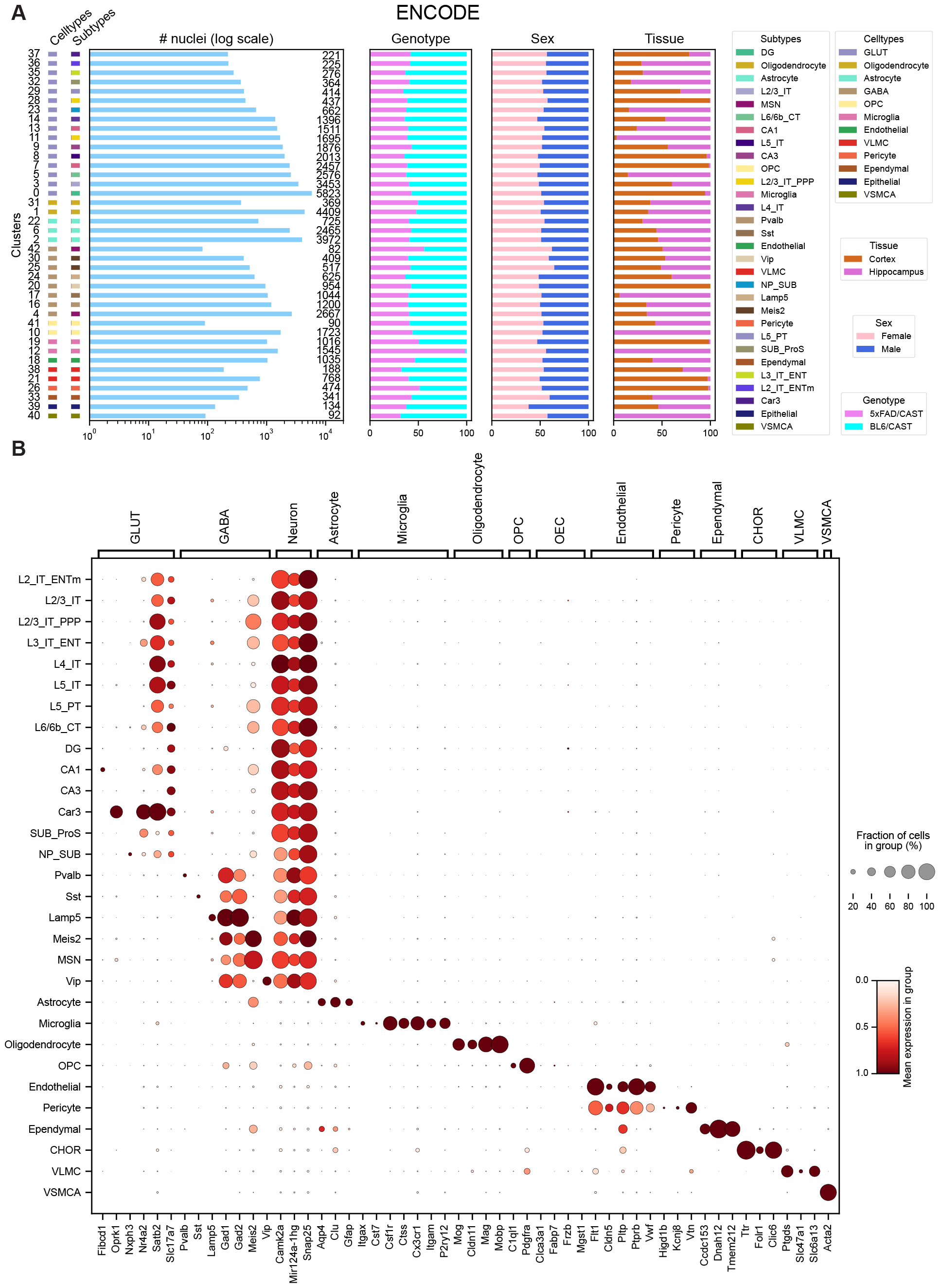
Overview of ENCODE dataset. **A**. Breakdown of nuclei, cell type, subtype, genotype, sex, and tissue across 43 clusters. **B**. Expression of marker genes

